# Sex-dependent adipose glucose partitioning by the mitochondrial pyruvate carrier

**DOI:** 10.1101/2024.05.11.593540

**Authors:** Christopher E Shannon, Terry Bakewell, Marcel J Fourcaudot, Iriscilla Ayala, Giovanna Romero, Mara Asmis, Leandro C Freitas Lima, Martina Wallace, Luke Norton

**Author notes:** **Corresponding author:** Christopher Shannon, PhD; Luke Norton, PhD.

## Abstract

**Objective:** The mitochondrial pyruvate carrier (MPC) occupies a critical node in intermediary metabolism, prompting interest in its utility as a therapeutic target for the treatment of obesity and cardiometabolic disease. Dysregulated nutrient metabolism in adipose tissue is a prominent feature of obesity pathophysiology, yet the functional role of adipose MPC has not been explored. We investigated whether the MPC shapes the adaptation of adipose tissue to dietary stress in female and male mice.

**Methods:** The impact of pharmacological and genetic disruption of the MPC on mitochondrial pathways of triglyceride assembly (lipogenesis and glyceroneogenesis) was assessed in 3T3L1 adipocytes and murine adipose explants, combined with analyses of adipose MPC expression in metabolically compromised humans. Whole-body and adipose-specific glucose metabolism were subsequently investigated in male and female mice lacking adipocyte MPC1 (*Mpc1*^AD-/-^) and fed either standard chow, high-fat western style, or high-sucrose lipid restricted diets for 24 weeks, using a combination of radiolabeled tracers and GC/MS metabolomics.

**Results:** Treatment with UK5099 or siMPC1 impaired the synthesis of lipids and glycerol-3-phosphate from pyruvate and blunted triglyceride accumulation in 3T3L1 adipocytes, whilst MPC expression in human adipose tissue was negatively correlated with indices of whole-body and adipose tissue metabolic dysfunction. Mature adipose explants from *Mpc1*^AD-/-^ mice were intrinsically incapable of incorporating pyruvate into triglycerides. *In vivo*, MPC deletion restricted the incorporation of circulating glucose into adipose triglycerides, but only in female mice fed a zero fat diet, and this associated with sex-specific reductions in tricarboxylic acid cycle pool sizes and compensatory transcriptional changes in lipogenic and glycerol metabolism pathways. However, whole-body adiposity and metabolic health were preserved in *Mpc1*^AD-/-^ mice regardless of sex, even under conditions of zero dietary fat.

**Conclusion:** These findings highlight the greater capacity for mitochondrially driven triglyceride assembly in adipose from female versus male mice and expose a reliance upon MPC-gated metabolism for glucose partitioning in female adipose under conditions of dietary lipid restriction.

## INTRODUCTION

Adipose tissue contributes to the intricate regulation of whole-body energy metabolism through the cyclic synthesis and breakdown of triglyceride stores. Nutrient fluxes into healthy adipocytes serve to buffer postprandial increases in circulating lipids and glucose whilst, in the postabsorptive state, lipolysis and free fatty acid (FFA) re-esterification balance systemic FFA availability with peripheral energy demands (Frayn, 2002; Virtanen et al., 2001). Under conditions of sustained nutrient excess, the progressive failure of these systems leads to chronic elevations in plasma FFA, ectopic lipid deposition (i.e. in skeletal muscle or liver) and the gradual worsening of peripheral insulin resistance (DeFronzo, 2004; Norton et al., 2022). Therefore, dysregulated nutrient handling in adipose tissue is an important hallmark of obesity that is directly implicated in the pathogenesis of common cardiometabolic diseases like type 2 diabetes and metabolic dysfunction-associated liver disease.

Adipose mitochondria are increasingly recognized as important modifiers of obesity-related pathophysiology. Unlike their counterparts in highly oxidative tissues (e.g. skeletal muscle and brown adipose tissue), mitochondria in white adipose tissue (WAT) are specialized for nutrient storage (Forner et al., 2009; Rotondo et al., 2017). Mitochondrial fluxes in WAT support triglyceride assembly by fueling the *de novo* synthesis of both fatty acids (lipogenesis; DNL) and glycerol-3-phosphate (glyceroneogenesis). Under normal dietary conditions, the contribution of DNL to adipose triglyceride storage in humans is believed to be less than 10% (Strawford et al., 2004). However, studies in rodents (Fernandez et al., 2019) and humans (Aarsland et al., 1997) have implicated an important role for adipose DNL in diverting dietary carbohydrates towards lipid storage, under conditions of excess carbohydrate availability. Moreover, impairments in adipose DNL and triglyceride storage have been associated with the development of insulin resistance in humans (Allister et al., 2015). Glyceroneogenesis is an important source of glycerol-3-phosphate generation in adipose (Nye et al., 2008) and facilitates FFA esterification to sustain the healthy deposition of triglycerides (Reshef et al., 2003). The regulation of these integrated mitochondrial pathways thus underpins the adaptive plasticity of WAT following dietary perturbations (Felix et al., 2021).

The mitochondrial pyruvate carrier (MPC) regulates cellular nutrient partitioning by gating pyruvate entry into mitochondria (Bricker et al., 2012; Herzig et al., 2012) where, in WAT, it is used both as a precursor for DNL and as the obligate substrate for glyceroneogenesis. Despite emergent tissue-specific functions for the MPC in numerous metabolic organs (Fernandez-Caggiano et al., 2020; Gray et al., 2015; Huang et al., 2016; McCommis et al., 2015; McCommis et al., 2016; McCommis et al., 2020; Panic et al., 2020; Shannon et al., 2017; Sharma et al., 2019; Wenes et al., 2022), the relevance of MPC activity in WAT has received surprisingly little attention. *In vitro* data demonstrate that mitochondrial pyruvate transport may be dispensable for the differentiation of 3T3L1 pre-adipocytes (Burrell et al., 2020), but this does not preclude a role for the MPC in mature WAT *in vivo*. On the contrary, prior studies suggest that reductions in WAT MPC expression, either by heterogenous knockdown (Zou et al., 2018) or in response to a high fat diet (Burrell et al., 2020), might be associated with alterations in systemic metabolism. However, the effects of MPC manipulation in WAT have not been directly investigated and thus its functional relevance in this tissue remains unclear.

Our understanding of the mechanisms through which dietary perturbations trigger adipose dysfunction has been convoluted by (often overlooked) sex-specific differences in adipose nutrient handling. For example, female adipose tissue displays greater intrinsic capacity for triglyceride synthesis and lipolysis than males, especially in subcutaneous depots (Edens et al., 1993; Fernandez et al., 2019). Moreover, sex differences in adipose tissue influence the phenotypic responses to different diets (Stanhope et al., 2009) and likely contribute to the increased incidence of obesity-related complications in males versus females (Després et al., 2000). Despite their fundamental importance in the development of precision therapeutics against obesity-related disease, the molecular events underpinning sex-specific responses to dietary stress remain poorly characterized.

We reasoned that mitochondrial substrate partitioning might shape the sex-specific impact of dietary nutrients on metabolic health and, specifically, that the MPC would play a central role in this process. To address these questions, we explored the relationship between MPC-gated intermediary metabolism and adipocyte function, assessed the requirement for MPC of metabolic adaptations to dietary lipid restriction and excess, and compared the role of WAT MPC between male and female mice.

## MATERIALS AND METHODS

### 2.1. Experimental Models and Animal Details

#### 2.1.1. Cell culture

3T3L1 adipocytes were maintained in DMEM supplemented with GlutaMAX^TM^ and were differentiated in absence of thiazolidinediones as previously described (Chen et al., 2018). For siRNA transfection experiments, 3T3L1 adipocytes were trypsin detached five days post-differentiation and reverse transfected with 50 pmols of Dharmacon SmartPool siSCR (#D-001810-10-05) or siMPC1 (#L-040908-01-0005) using Lipofectamine® RNAiMAX reagent (#13778; ThermoFisher, MA, USA) following published protocols (Kilroy et al., 2009). Based upon initial optimization experiments of MPC1 depletion (Supplemental Figure 1A), subsequent assays were performed 5 days post transfection.

#### 2.1.2. Human Studies

All studies involving human subjects were approved by the University of Texas Health Science Center at San Antonio Institutional Review Board. Fasting abdominal subcutaneous adipose tissue specimens were obtained from subjects with and without adipose tissue insulin resistance (AdipoIR), as previously described (Winnier et al., 2015). All subjects underwent a standard 2-h oral GTT to confirm normal glucose tolerance (NGT) or impaired glucose tolerance (IGT). Fasting AdipoIR was calculated from free fatty acid and insulin data (Gastaldelli et al., 2017).

#### 2.1.3. Animal Studies

Mice with LoxP sites flanking the Mpc1 allele, backcrossed through C57Bl/6J mice, were a gift from Eric Taylor and have been described previously (Gray et al., 2015). Adiponectin-Cre (Adipoq-Cre) mice were purchased from The Jackson Laboratory (stock # 010803) (Bar Harbor, ME). LoxP/LoxP littermate controls were used in all experiments. Mice were housed in environmentally controlled conditions (23°C, 12-hour light/dark cycles), provided ad-libitum access to water and a fed a standard chow or, starting at 6 weeks of age, a zero fat diet (D04112303) or a western-style high fat diet (D09100310) from Research Diets Inc. (NJ, USA). Experiments were conducted in 32-week aged mice following 24-26 weeks of diet treatments. All procedures were approved by the Institutional Animal Care and Use Committee at University of Texas Health Science Center at San Antonio.

### 2.2. Method Details

#### 2.2.1. Western blotting and quantitative real-time polymerase chain reaction (qRT-PCR)

Immunoblot analysis was carried out on cell or adipose lysates using mouse or human primary antibodies against MPC1 (Sigma #HPA045119), MPC2 (Cell Signaling Technologies #46141), GAPDH (Sigma #G8795) and β-Tubulin (Abcam #179513) and developed by chemiluminescence (ECL). QRT-PCR was performed using pre-designed TaqMan probes (Life Technologies, CA, USA) or SYBR green primer assays (Integrated DNA Technologies, Iowa, USA). Data were normalized to the geometric mean of the reference genes *B2m, Hmbs, Gapdh, Rplpo*, and *β-actin*.

#### 2.2.2. [2,3-^14^C] pyruvate incorporation assays

Approximately 5×10^5^ adherent 3T3L1 adipocytes, eight days post-differentiation, were washed twice with PBS and incubated with complete growth media supplemented with 0.1 µCi/ml [2,3-^14^C] pyruvate or [U-^14^C] acetate with or without UK5099 (5uM) or insulin (300 pM) for six hours. Cells were subsequently washed with PBS and quenched with ice cold methanol. For explant studies, 20-30 mg of freshly excised WAT was incubated in media as above, then washed with PBS and snap frozen in liquid nitrogen. Total lipids were extracted from cell lysates or frozen extracts by the Folch method (FOLCH et al., 1957) and dried under oxygen-free nitrogen. Lipid residues were saponified in methanolic KOH (1.5M) for one hour at 70°, neutralized with methanolic HCl (3M) and re-extracted for separation of glycerol-glyceride and fatty acyl fractions. Dried fractions were resuspended and mixed with scintillation fluid prior to scintillation counting.

#### 2.2.3. Triglyceride content

Triglyceride content was determined in 3T3L1 lysates (5% NP-40) using a calorimetric kit (Ab65336, Abcam) following the manufacturer’s instructions.

#### 2.2.4. *Ex vivo* lipolysis assays

Glycerol and free fatty acid release from freshly excised inguinal and epididymal fat explants (20-30 mg) were assessed under basal, insulin-treated, and maximal (10 µM forskolin / 5 µM Triacsin C) conditions using published protocols (Baskaran & Thyagarajan, 2017). Media glycerol (F6428, Sigma) and NEFA (Wako) concentrations were determined by calorimetric assay.

#### 2.2.5. Mouse physiology studies

##### 2.2.5.1. Body Composition

Lean mass, fat mass and total body water were determined in 24-week-old WD and ZFD mice by qMRI at the San Antonio Nathan Shock Centre Aging Animal Models and Longevity Assessment Core.

##### 2.2.5.2. Oral Glucose Tolerance Test

Overnight fasted (16 hours) mice were administered with dextrose (2 g/kg bodyweight) by oral gavage. Glucose concentration was determined in whole-blood sampled at baseline and at 15-, 30-, 60- and 120-minute intervals from a tail incision using a glucometer (Contour next EZ).

##### 2.2.5.3. [U-^14^C] Glucose incorporation Assays

Overnight fasted mice were administered 2 nCi [U-^14^C] glucose in 500 µl saline by intraperitoneal injection and were euthanized by isoflurane (inhaled) five hours later. Tissues were excised, snap frozen in liquid nitrogen and stored at -80°, with ∼200 mg portions used for lipid extraction and scintillation counting as described above.

#### 2.2.6. Plasma Analytes

Plasma insulin was determined by ELISA (#90080; Crystal Chem, IL, USA). Plasma free fatty acids (999-34691; FUJIFILM Wako Chemicals; VA, USA), glycerol (F6428; Sigma, MO, USA) and triglycerides (TR22421; ThermoFisher, MA, USA) were determined by enzymatic calorimetric assay.

#### 2.2.7. GC/MS Polar Metabolomics

Approximately 50 mg of tissue plus 30 pmol/mg tissue of internal standard (MSK-A2-1.2, Cambridge Isotope Laboratories, Inc) were extracted in chloroform:methanol:saline (2:1:1) by bead homogenization. Extracts were centrifuged at 10,000g for 5 min at 4°C and the upper layer (polar fraction) was evaporated to dryness under oxygen-free nitrogen. Residues were reconstituted in 20 µl methoxyamine (20 mg/ml in pyridine) and incubated at 37°C for one hour. 20 µl MBSTFA was added prior to a further 30-minute incubation at 37°C. Polar derivatives were analysed by GC–MS using a DB-35MS column (30 m × 0.25 mm internal diameter × 0.25 μm, Agilent J&W Scientific) installed in an Agilent 7890 A gas chromatograph interfaced with an Agilent 5975 C mass spectrometer as previously described (Cordes & Metallo, 2019). Spectra were analysed in MatLab and natural isotope abundance was corrected using a custom-built script (Fernandez et al., 1996). Metabolite abundances were normalized to their respective internal standard (in pmol / mg tissue weight) or, for metabolites without a respective internal standard, normalized to ^13^C valine (relative abundance).

### 2.3. Quantification and statistical analyses

Differences between genotypes, sexes and/or diets were assessed using two-way analysis of variance with multiple comparisons controlled post-hoc, as appropriate. Further details for individual comparisons are provided in figure legends. Significance testing was performed using GraphPad Prism 10. Data are presented as mean ± standard error, and data were considered significantly different at P < 0.05.

## RESULTS

### Mitochondrial pyruvate transport maintains triglyceride storage in 3T3L1 adipocytes

Proteomic screening recently identified the mitochondrial pyruvate carriers MPC1 and MPC2 as two of the most highly induced proteins during adipocyte differentiation (Choi et al., 2020). To further probe the functional significance of mitochondrial pyruvate transport in adipocytes, we next monitored [2-^14^C] pyruvate incorporation into cellular lipids. Treatment of differentiated adipocytes cultured in complete growth media with the MPC inhibitor UK5099 (5 µM) led to a ∼50% reduction in pyruvate incorporation into the lipid pool (Figure 1A). However, near-complete suppression of pyruvate-driven lipid synthesis was observed when adipocytes were treated with UK5099 in serum-free media (Figure 1A). Pyruvate flux into lipids was stimulated by insulin and was partially dependent upon both glucose and glutamine availability but was abrogated by UK5099 treatment regardless of substrate conditions (Figure 1A).

**Figure 1:**
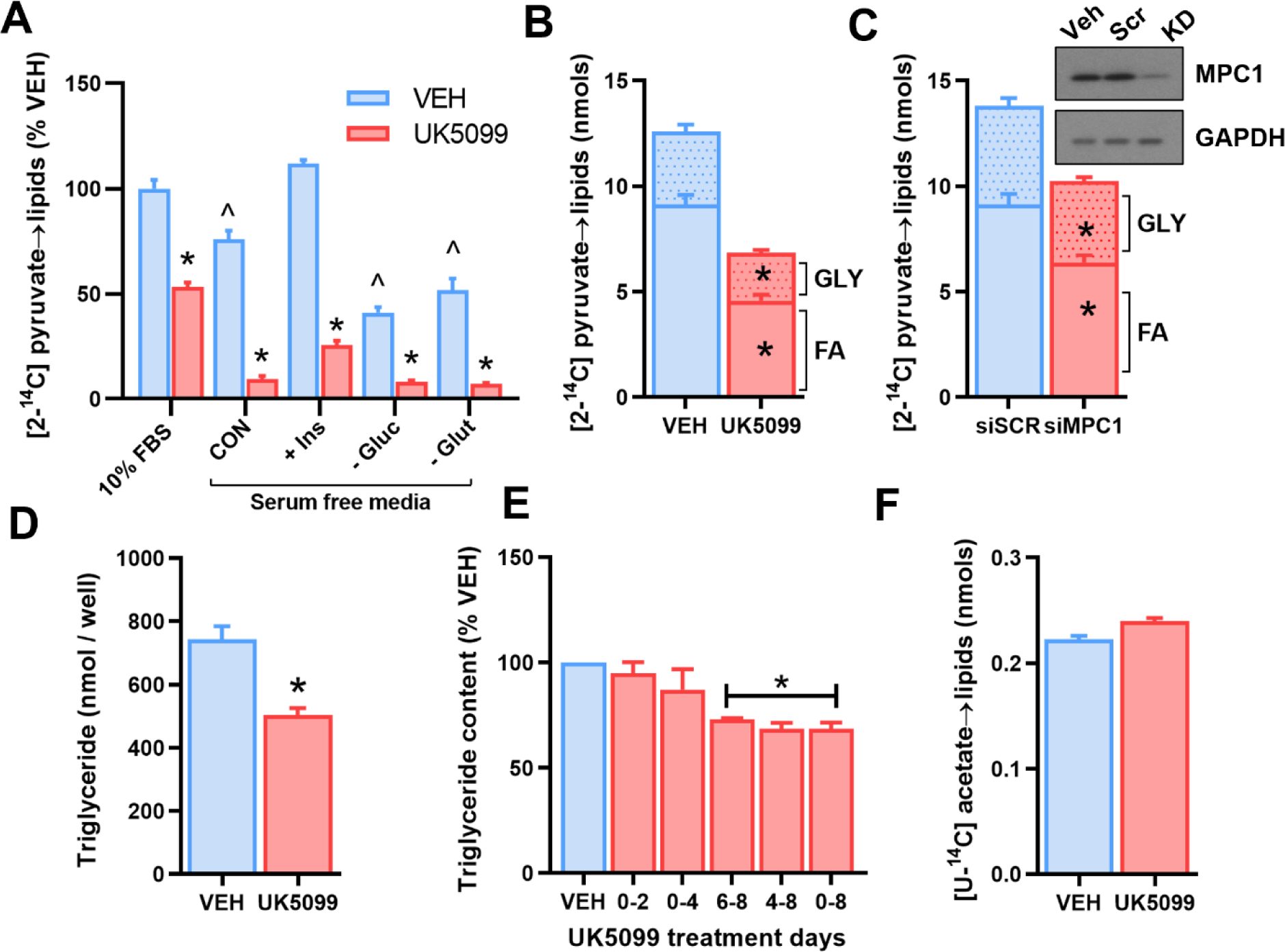
Mitochondrial pyruvate transport maintains triglyceride storage in 3T3L1 adipocytes. **(A - C)** Incorporation of [2-^14^C] pyruvate into total lipids (**A**), or the fatty acyl or glycerol-glyceride fraction of lipids (**B – C**) following treatment of differentiated 3T3L1 adipocytes with 5µM UK5099 **(A – B)** or siRNA against MPC1 or scramble control **(C)**. **(D - E)** Total triglyceride accumulation in adipocytes treated with 5µM UK5099 either throughout **(D)**, or at various time points during **(E)**, differentiation. **(F)** Incorporation of [U-^14^C] acetate into total lipids in differentiated 3T3L1 adipocytes treated with or without 5µM UK5099. Data are mean ± SE for at least three experimental replicates (different cell passages). *P<0.05 vs control, ^P<0.05 vs 10% FBS condition in **(A)** by paired t-test (**B** - **D** and **F**), one-way ANOVA (**E**) or 2-way ANOVA (**A**).

Incorporation of pyruvate into the total lipid pool reflects the *de novo* synthesis (and subsequent esterification) of either fatty acids or glycerol-3-phosphate (gly-3P). Tracing pyruvate into the fatty acyl or gly-3P moieties of total lipids demonstrated that both these pathways were sensitive to inhibition by UK5099 (Figure 1B). The reliance of pyruvate-driven lipogenesis on MPC-linked metabolism was further validated in 3T3-L1 adipocytes transfected with *Mpc1* siRNA to deplete MPC abundance by ∼50% (Figure 1C and Supplementary Fig 1). Since precursors other than pyruvate (e.g. glutamine) could support anabolic pathways in adipocytes (Roberts et al., 2009), we next assessed the overall dependence of triglyceride accumulation on mitochondrial pyruvate transport. Treatment with UK5099 during adipocyte differentiation, despite the presence of 2 mM glutamine, resulted in a 35% reduction in total triglyceride content (Figure 1F). Notably, delaying UK5099 treatment until the end of differentiation (days 6-8) suppressed triglyceride levels to a similar extent as to when UK5099 treatment was maintained throughout differentiation (Figure 1E), whereas treatment on days 0-4 had a negligible impact. Importantly, lipogenic pathways downstream of the MPC were unaffected by UK5099 treatment, as shown by the preservation of [U-^14^C] acetate incorporation into the triglyceride pool (Figure 1F). These data extend previous observations that the MPC is dispensable for the early events of adipogenesis (Burrell et al., 2020), illustrating an obligate role for pyruvate-driven metabolism in maintaining triglyceride accumulation in differentiated adipocytes.

### Adipose tissue MPC expression is altered in metabolically compromised mice and humans

The ability to accumulate triglycerides is an important function of healthy adipose tissue and this becomes dysregulated under metabolic stressors such as obesity and type 2 diabetes (Palavicini et al., 2021; Pietiläinen et al., 2011). To establish the pathological relevance of mitochondrial pyruvate transport in mature adipose tissue, we quantified adipose MPC expression in tissue from metabolically compromised mice and humans. Both MPC1 and MPC2 proteins were significantly lower in the subcutaneous inguinal WAT (iWAT) from obese versus lean mice (Figure 2A). Similarly, MPC1 and MPC2 protein expression was reduced in subcutaneous WAT from prediabetic human subjects displaying impaired fasting glucose, impaired glucose tolerance, and impaired adipose function compared to control subjects with normal glucose tolerance (Figure 2B and Supplementary Fig 2). Moreover, adipose tissue MPC expression was negatively correlated with oral glucose tolerance (Figure 2C-D and F-G) and adipose tissue insulin resistance (Figures 2E and H) across all subjects. Together with our *in vitro* experiments, these findings highlight a potential link between reduced MPC expression in WAT, adipose tissue function, and the progression of metabolic disease.

**Figure 2:**
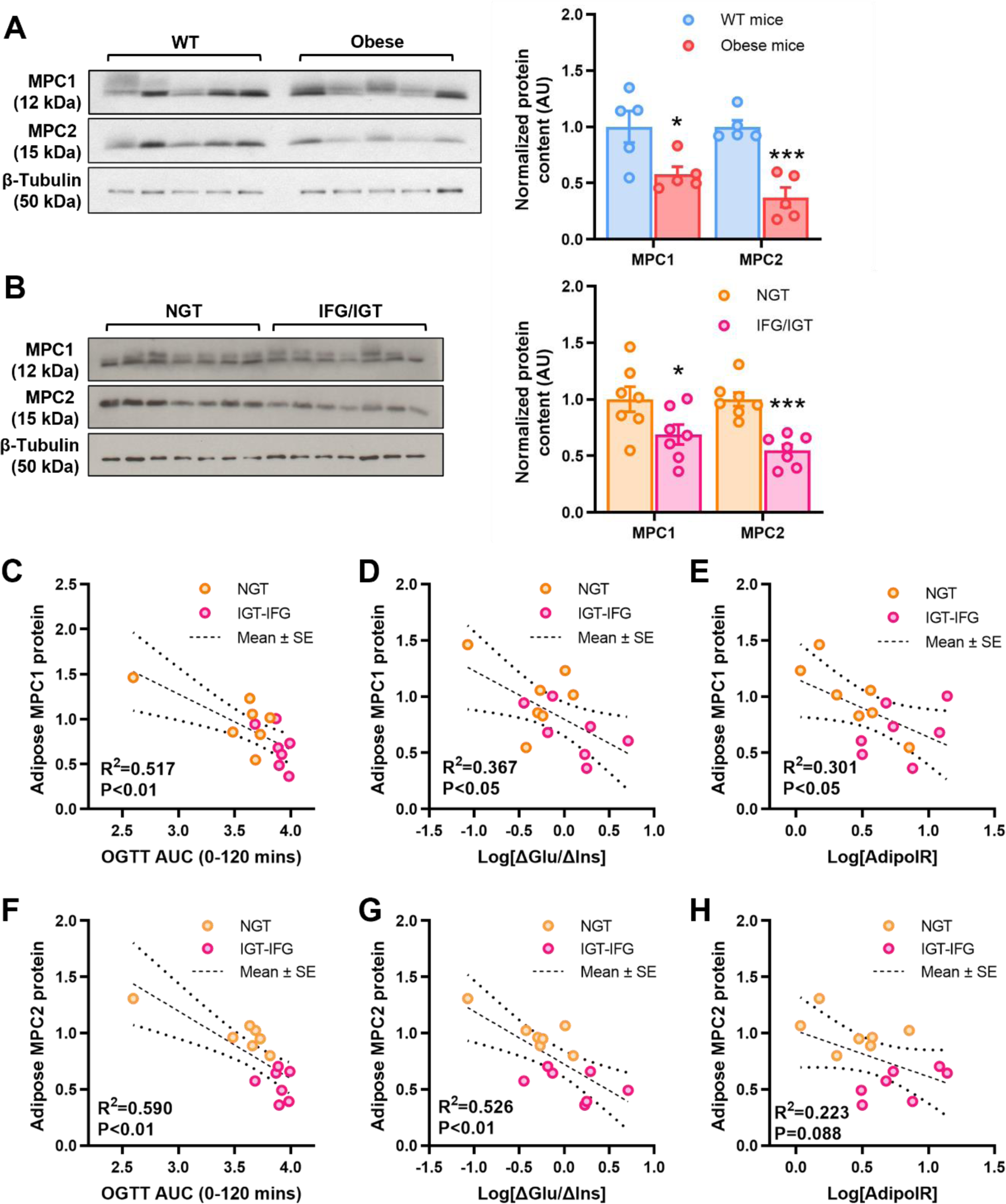
Adipose tissue MPC expression is altered in metabolically compromised mice and humans. **(A - B)** Normalized, quantified protein expression and representative western blot of mitochondrial pyruvate carrier proteins in subcutaneous adipose tissue from wild type vs genetically obese (ob/ob) mice (**A**) and from human subjects with normal glucose tolerance vs impaired fasting glucose plus impaired glucose tolerance (**B**). (**C** - **H**) Pearson correlations between the protein expression of MPC1 (**A**, **D**, **E**) or MPC2 (**F**, **G**, **H**) in subcutaneous adipose tissue and indices of glucose tolerance (**C** and **F**), insulin sensitivity (**D** and **G**) and adipose insulin resistance (**E** and **H**) in human subjects. Data are n=5 (**A**) or n=7 (**B** – **H**) in each group. *P<0.05, ***P<0.001 vs control group (**A** and **B**).

### MPC1 knockdown blunts capacity for triglyceride synthesis in mature adipose tissue

To test the hypothesis that a reduction in adipose tissue MPC expression could exacerbate metabolic dysfunction by restricting the healthy expansion of triglyceride stores, adipose specific *Mpc1* knockout mice were generated using the Cre-lox system with an Adiponectin-Cre promoter. Cre recombination in homozygous floxed mice silenced MPC1 protein expression in subcutaneous adipose tissue from *Mpc1*^AD-/-^ mice (Figure 3A). To examine the functional significance of adipose MPC ablation, we first compared *ex vivo* rates of pyruvate-driven lipid synthesis in adipose tissue explants from *Mpc1*^AD-/-^ mice with LoxP^+/+^ controls. Basal and insulin-stimulated rates of lipogenesis were markedly suppressed in visceral epididymal (eWAT, Figure 3B) and iWAT (Supplemental Fig 3A) explants from *Mpc1*^AD-/-^ mice. Moreover, residual rates of lipogenesis in *Mpc1*^AD-/-^ explants were insensitive to further inhibition by UK5099 (Figure 3B), confirming the complete loss of MPC gated lipid synthesis in *Mpc1*-silenced adipose tissue. In agreement with our *in vitro* findings in 3T3-L1 cells, pyruvate-driven gly-3P synthesis was also compromised, by 50-90%, in *Mpc1*^AD-/-^ explants (Figure 3C). Notably, rates of both lipogenesis and gly-3P synthesis were substantially higher in explants from female mice compared to males (Figure 3B and C and Supplemental Fig 3A), which is consistent with previous reports on sexual dimorphism in the lipogenic capacity of adipose tissue (Fernandez et al., 2019; Macotela et al., 2009).

**Figure 3:**
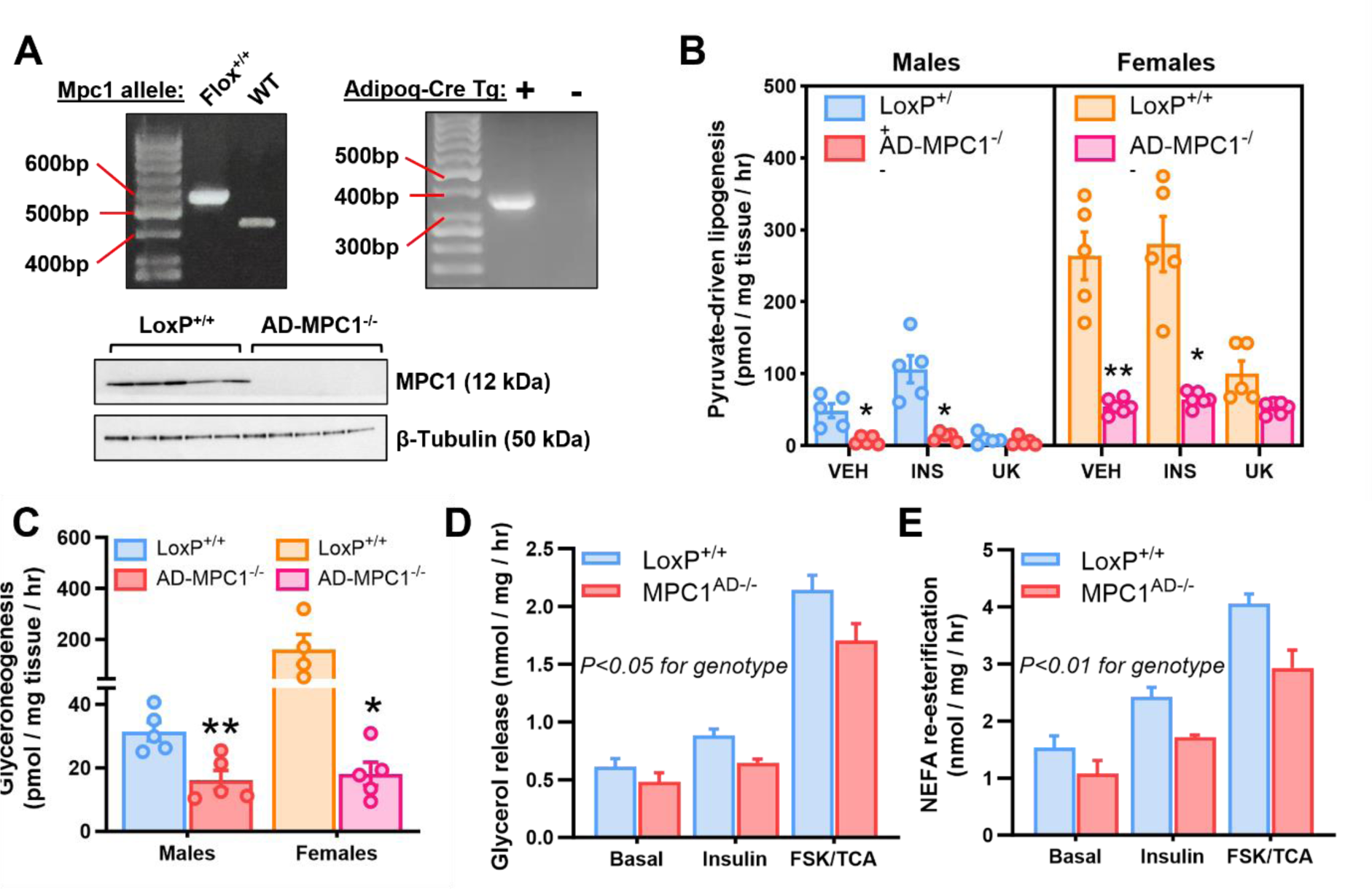
MPC1 knockdown blunts capacity for triglyceride synthesis in mature adipose tissue. **(A)** MPC1 floxed allele and Adipoq-Cre recombination was visualized by semiquantitative RT-PCR and MPC1 protein knockdown efficacy was verified by western blot of adipose lysates. (**B - C**) *Ex vivo* incorporation of [2-^14^C] pyruvate into the fatty acyl (lipogenesis **B**) or glycerol (glyceroneogenesis **C**) moieties of total lipids in epididymal adipose explants from male and female MPC^AD-/-^ mice or LoxP^+/+^ controls. (**D** – **E**) Rates of glycerol release (**D**) and non-esterified fatty acid re-esterification (**E**) from epididymal adipose explants from male MPC^AD-/-^ mice or LoxP^+/+^ controls under basal (unstimulated) conditions or during treatment with insulin or forskolin plus triacsin C. *P<0.05, **P<0.01 for MPC^AD-/-^ vs LoxP^+/+^; ^^P<0.01 for female vs male. Data are mean ± SE for at least five mice per group.

A considerable fraction of the FFA liberated from adipose TAG during lipolysis is re-esterified, requiring a readily available pool of gly-3P (Nye et al., 2008; Reshef et al., 2003). We therefore investigated whether the impairment in pyruvate-driven gly-3P synthesis due to MPC deletion also impacted upon the capacity for lipolysis or FFA recycling in WAT. Free glycerol release from iWAT explants was slightly lower in *Mpc1*^AD-/-^ mice compared to LoxP^+/+^ controls (Figure 3D), with the greatest effect observed when lipolysis was stimulated with forskolin and triacsin C. A similar trend was observed in eWAT (Supplemental Fig 3B), indicating a blunted lipolytic capacity in *Mpc1*^AD-/-^ WAT. Moreover, this reduction in lipolysis was paralleled by lower rates of FFA re-esterification in *Mpc1*^AD-/-^ explants (Figure 2E and Supplemental Fig 3C). As a result of the similar reduction in both lipolysis and FFA re-esterification, absolute rates of explant NEFA release were comparable between *Mpc1*^AD-/-^ and LoxP^+/+^ mice (Supplemental Fig 3D-E). Thus, MPC1 knockout compromises the intrinsic capacity of mature adipose tissue for both anabolic (glyceroneogenesis, lipogenesis and FFA recycling) and catabolic (lipolysis) processes relevant to metabolic function.

### Sex-specific dependencies on the MPC for adipose DNL in vivo

Circulating glucose is a major precursor of the intracellular pyruvate pool and an important physiological substrate for the subsequent synthesis of glycerol-3-phosphate and fatty acids in adipose. Accordingly, we next determined whether loss of MPC-gated lipid synthesis impacted upon the incorporation of an oral bolus of [U-^14^C] glucose into adipose lipids in Mpc1^AD-/-^ and LoxP^+/+^ mice. Since reliance on adipose DNL may be augmented under certain dietary conditions (Fernandez et al., 2019), mice were challenged with either a high-fat western-style diet (WD) or a zero-fat sucrose-enriched diet (ZFD) for 24 weeks.

In iWAT, the conversion of glucose into fatty acids (i.e. DNL from glucose) was significantly higher in LoxP^+/+^ females compared to males and was robustly increased by ZFD versus WD in both sexes (Figure 4A). Whereas the induction of DNL with ZFD was preserved in iWAT from *Mpc1*^AD-/-^ males, it was blunted by 43% (Figure 4A) in *Mpc1*^AD-/-^ females. In eWAT, DNL from glucose was unaffected by diet or genotype for male mice but was again markedly reduced in *Mpc1*^AD-/-^ females fed ZFD (Figure 4B). These data show that reliance on MPC-gated metabolism for glucose flux into fatty acids is greater in female than male adipose.

**Figure 4:**
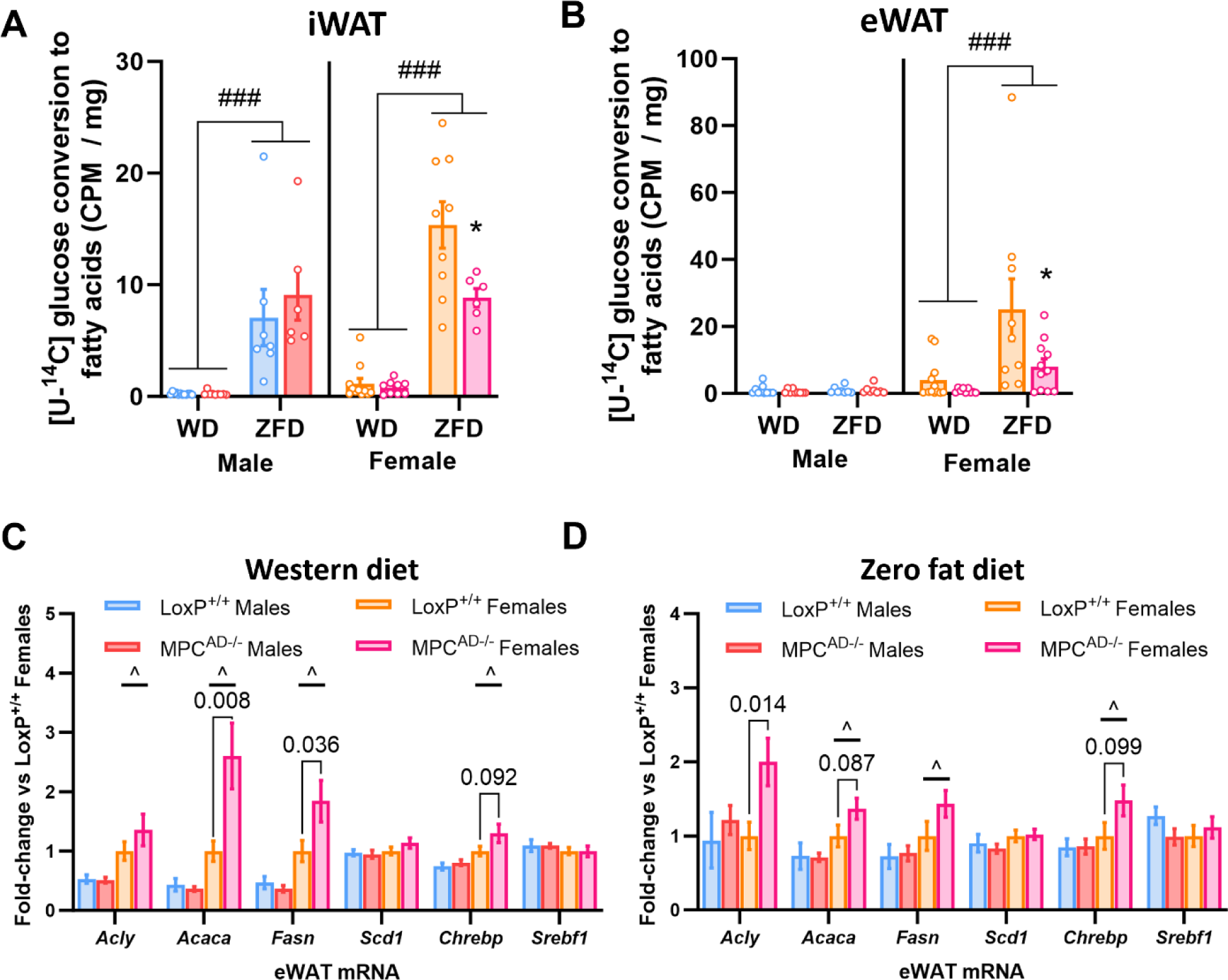
Whole body lipid and glucose homeostasis are preserved in chow fed Mpc1^AD-/-^ mice. (**A**) Total body weight, (**B**) inguinal and epididymal adipose depot weights, (**C**) oral glucose tolerance, (**D**) HOMA-IR index of insulin resistance, (**E - F**) fasting and CL316,243-stimulated plasma glycerol (**E**) and non-esterified fatty acids (**F**) in male and female MPC^AD-/-^ mice or LoxP^+/+^ controls fed a standard chow diet. *P<0.05 for MPC^AD-/-^ vs LoxP^+/+^; ^^^P<0.001 for female vs male. Data are mean ± SE for 8-10 mice per group.

To gain further insight into the rewiring of adipose DNL in *Mpc1*^AD-/-^ females, we also measured the mRNA expression of lipogenic enzymes and transcription factors. Consistent with our *ex vivo* and *in vivo* tracer data, lipogenic gene expression was typically higher in adipose depots from female mice compared to males, especially in eWAT (Figure 4C-D and Supplemental Fig. 4A and B). Moreover, certain lipogenic genes, including *Acly*, *Acaca* and *Fasn*, were upregulated in adipose depots from female, but not male, *Mpc1*^AD-/-^ mice (Figure 4C and D and Supplemental Fig. 4A and B). These data suggest that transcriptional upregulation of rate-limiting lipogenic enzymes downstream of the MPC might compensate for the loss of glucose dependent DNL flux in MPC-deficient adipose tissue, especially in female mice.

Blockade of adipose DNL has previously been shown to promote a compensatory increase in liver DNL under ZFD conditions (Fernandez et al., 2019). As with adipose tissue, incorporation of [U-^14^C] glucose into hepatic fatty acids was greater following ZFD than WD, especially in female mice, but no differences were detected between *Mpc1*^AD-/-^ and LoxP^+/+^ mice regardless of sex (Supplemental Fig. 4C). Similarly, lipogenic gene expression, although increased in female livers versus males, was unchanged by the loss of adipose MPC (Supplemental Fig. 4D).

### Sex-specific dependencies on the MPC for adipose glycerol-3-phosphate synthesis in vivo

We next monitored the incorporation of [U-^14^C] glucose into the Gly-3P moiety of adipose lipids, which captures MPC-dependent glyceroneogenesis following the conversion of glucose to pyruvate, as well as the MPC-independent direct synthesis of Gly-3P from glucose via glycolysis. In iWAT, Gly-3P synthesis was higher in females than males, but largely unaffected by diet or genotype in either sex (Figure 5A). In contrast, whereas eWAT Gly-3P synthesis was again comparable between diets and genotypes in male mice, it was markedly upregulated by ZFD in LoxP^+/+^ females but entirely blunted in *Mpc1*^AD-/-^ females (Figure 5B).

**Figure 5:**
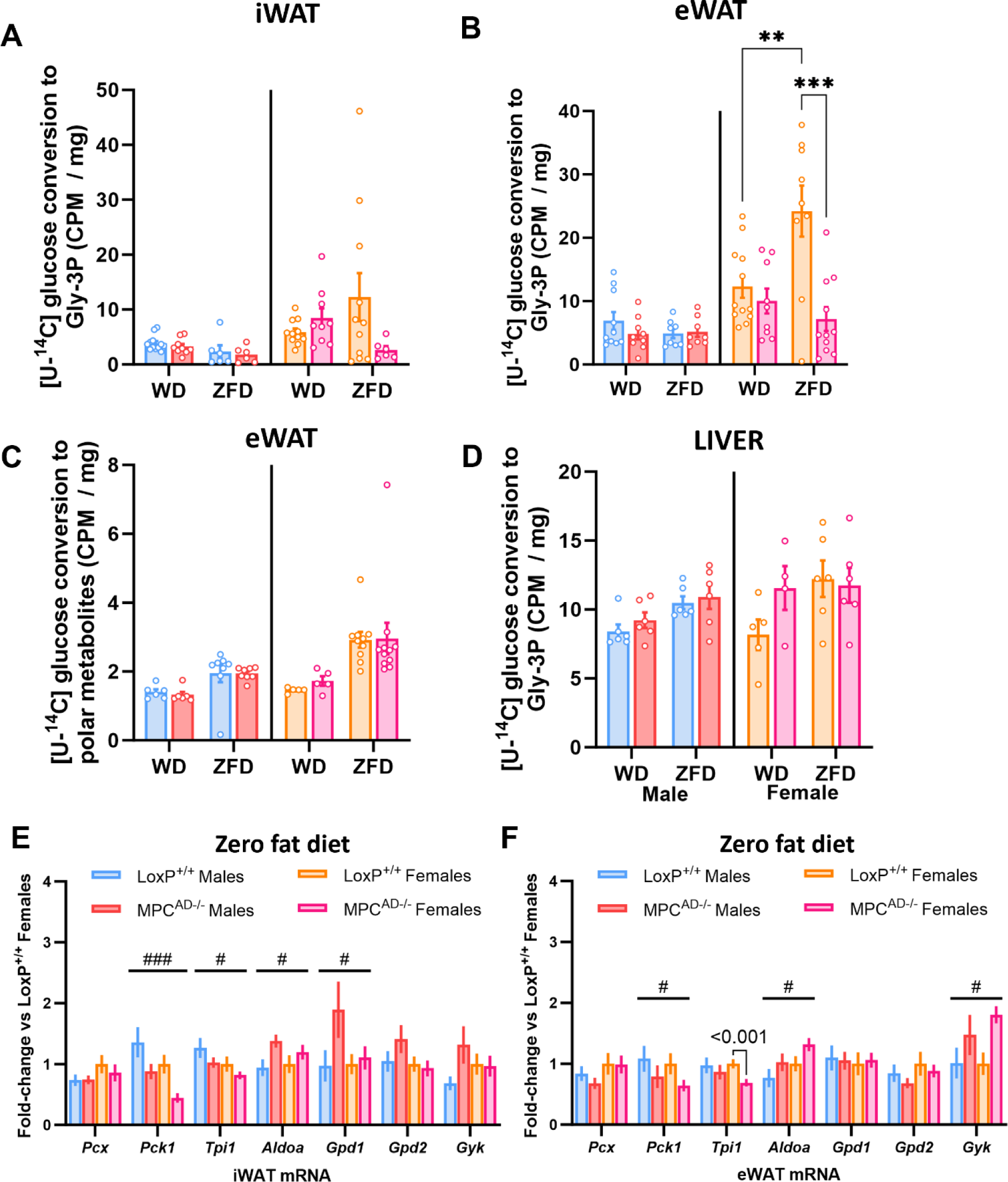
Total adiposity is maintained in Mpc1AD-/- mice fed a lipid restricted diet. (**A, D**) Percentage body weight gain, (**B**, **E**) total fat and lean mass, and (**C**, **F**) incremental oral glucose tolerance for male and female MPC^AD-/-^ mice or LoxP^+/+^ controls fed either a zero fat, sucrose enriched diet (**A** – **C**) or a high fat western-style diet (**D** – **F**) for 24 weeks. *P<0.05 for MPC^AD-/-^ vs LoxP^+/+^. Data are mean ± SE for 8-10 mice per group.

To confirm that the observed changes in gly-3P synthesis were not related to differences in adipose tissue glucose uptake, we also measured tracer enrichment in the polar fraction. Although flux from [U-^14^C] glucose into polar metabolites was increased by ZFD, and to a greater extent in female mice, it was unaffected by MPC knockdown (Figure 5C). Similarly, glucose incorporation into hepatic gly-3P was comparable between LoxP^+/+^ and *Mpc1*^AD-/-^ mice, regardless of sex or diet (Figure 5D). Thus, glucose trafficking towards gly-3P in visceral adipose is increased under conditions of dietary lipid restriction, but only in female mice and, moreover, this adaptation is entirely reliant upon MPC-mediated glyceroneogenesis.

We also investigated whether the transcriptional regulation of glyceroneogenesis was altered by MPC knockdown. Notably, *Pck1*, which encodes for the rate-limiting enzyme glyceroneogenic enzyme pyruvate carboxykinase, was consistently reduced in adipose depots from ZFD fed *Mpc1*^AD-/-^ mice, regardless of sex (Figure 5E-F). We also observed changes in *Aldoa* (increased) and *Tpi1* (decreased), both terminal enzymes in the glycolytic generation of glyceraldehyde-3-phosphate, as well as increased *Gdp1*, responsible for the final step of gly-3P synthesis, in *Mpc1*^AD-/-^ adipose. Interestingly, glycerol kinase (Gyk) expression was increased in *Mpc1*^AD-/-^ iWAT (Figure 5F), indicating a potential role for free glycerol as an alternative substrate for Gly-3P synthesis via triglyceride-glycerol recycling (Guan et al., 2002).

### Decreased adipose TCA cycle pool size in female Mpc1^AD-/-^ mice fed a lipid restricted diet

To better understand the metabolic pathways underpinning female-specific reliance on the MPC for DNL and gly-3P synthesis, we performed a targeted analysis of intermediary metabolites in eWAT from ZFD fed mice. We initially focused on intermediates within the TCA cycle, since both DNL and glyceroneogenesis rely on the anaplerotic and cataplerotic pathways of the TCA cycle. The relative pool size of TCA cycle intermediates in female LoxP^+/+^ adipose was double that of males but was 35% lower in *Mpc1*^AD-/-^ females (Figure 6A). This was largely driven by reductions in malate and succinate abundance in *Mpc1*^AD-/-^ females, although other TCA intermediates followed a similar pattern (Figure 6A). Thus, MPC deficiency disrupts the balance between anaplerosis and cataplerosis in adipose from *Mpc1*^AD-/-^ female, but not male mice, consistent with the sex-specific impairment of adipose DNL and gly-3P synthesis.

**Figure 6:**
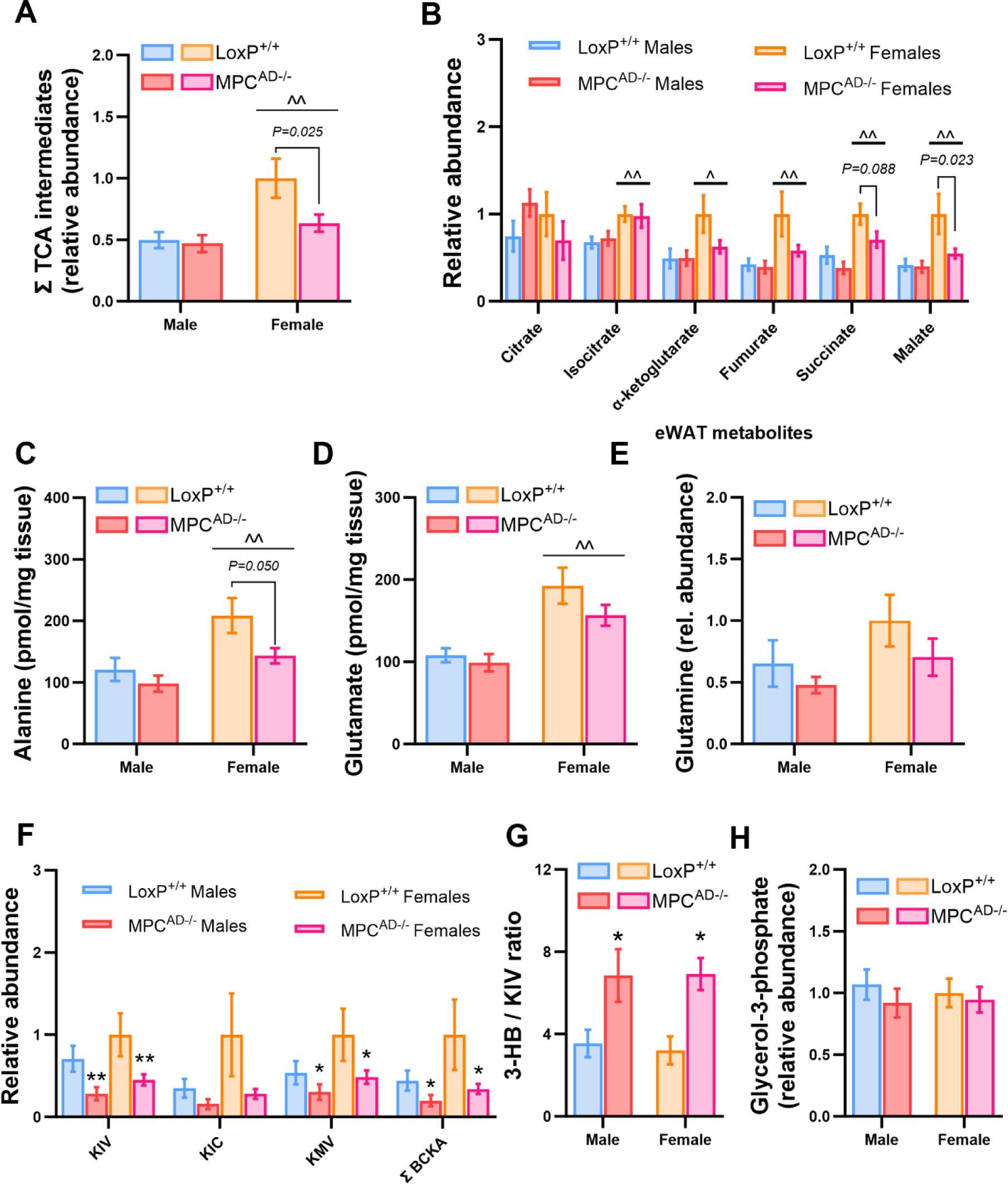
Sex-specific dependencies on the MPC for adipose DNL in vivo. (**A - B**) *In vivo* incorporation of intraperitoneally administered [U-^14^C] glucose into the fatty acyl moieties of total lipids (lipogenesis) in inguinal (**A**) and epididymal (**B**) adipose tissue depots and (**C** – **D**) mRNA expression of lipogenic genes in epididymal adipose tissue (eWAT) for male and female MPC^AD-/-^ mice or LoxP^+/+^ controls fed either a high fat western-style diet (WD, **C**) or a zero fat, sucrose enriched diet (ZFD, **D**) for 24 weeks. *P<0.05 for MPC^AD-/-^ vs LoxP^+/+^; ^###^P<0.001 for ZFD vs WD; ^P<0.05 for female vs male. Data are mean ± SE for 8-10 mice per group.

In absence of MPC-gated metabolism, alternate anaplerotic substrates (e.g. amino acids) have been shown to support TCA cycle dependent pathways (Gray et al., 2015; McCommis et al., 2015). Alanine levels were lower in adipose tissue from *Mpc1*^AD-/-^ mice, especially in females (Figure 6C), whilst glutamate (Figure 8D) and glutamine (Figure 6E) levels were not significantly different. Interestingly, branched chain keto acid levels were consistently reduced in *Mpc1*^AD-/-^ mice, regardless of sex (Figure 6E). Moreover, the ratio of 3-hydroxybutyrate to alpha keto isovalerate, a surrogate for branched chain keto dehydrogenase activity, was robustly increased in *Mpc1*^AD-/-^ mice (Figure 6F). These changes suggest that flux through branched chain amino acid (BCAA) catabolism is increased in MPC null adipose tissue and are consistent with recent reports following MPC inhibition in hepatocytes (Ferguson et al., 2023). Despite reductions in glucose driven gly-3P synthesis in *Mpc1*^AD-/-^ females (Figure 5B), levels of adipose gly-3P were comparable between *Mpc1*^AD-/-^ mice and LoxP^+/+^ controls (Figure 6H).

### Whole body lipid and glucose homeostasis are preserved in Mpc1^AD-/-^ mice fed a chow diet

Finally, we determined whether the observed alterations in adipose tissue lipid metabolism impacted weight gain and glucose homeostasis. Despite the observed *ex vivo* deficit in adipose triglyceride synthesis, weight gain was comparable between *Mpc1*^AD-/-^ and LoxP^+/+^ male mice and was marginally (∼5%) increased in *Mpc1*^AD-/-^ females, when mice were fed a standard chow diet (Figure 7A). Similarly, adipose depot weights were unchanged in *Mpc1*^AD-/-^ males, whereas iWAT mass was slightly greater in *Mpc1*^AD-/-^ females compared to LoxP^+/+^ controls (Figure 7B). No differences were found in glucose tolerance (Figure 7C) or HOMA-IR (fasting glucose x insulin; Figure 7D) between genotypes in either sex. Moreover, circulating free glycerol (Figure 7E) and non-esterified fatty acids (Figure 7F) were unaffected by the loss of adipose MPC1, either in the fasting state or upon simulation with the β_3_-adrenergic antagonist CL316,243 (10 mg/kg intraperitoneal injection). Together these data demonstrate that, under standard (chow) dietary conditions and regardless of biological sex, mitochondrial pyruvate transport in adipose is not required for the maintenance of whole-body glycemic or lipolytic control *in vivo*.

**Figure 7:**
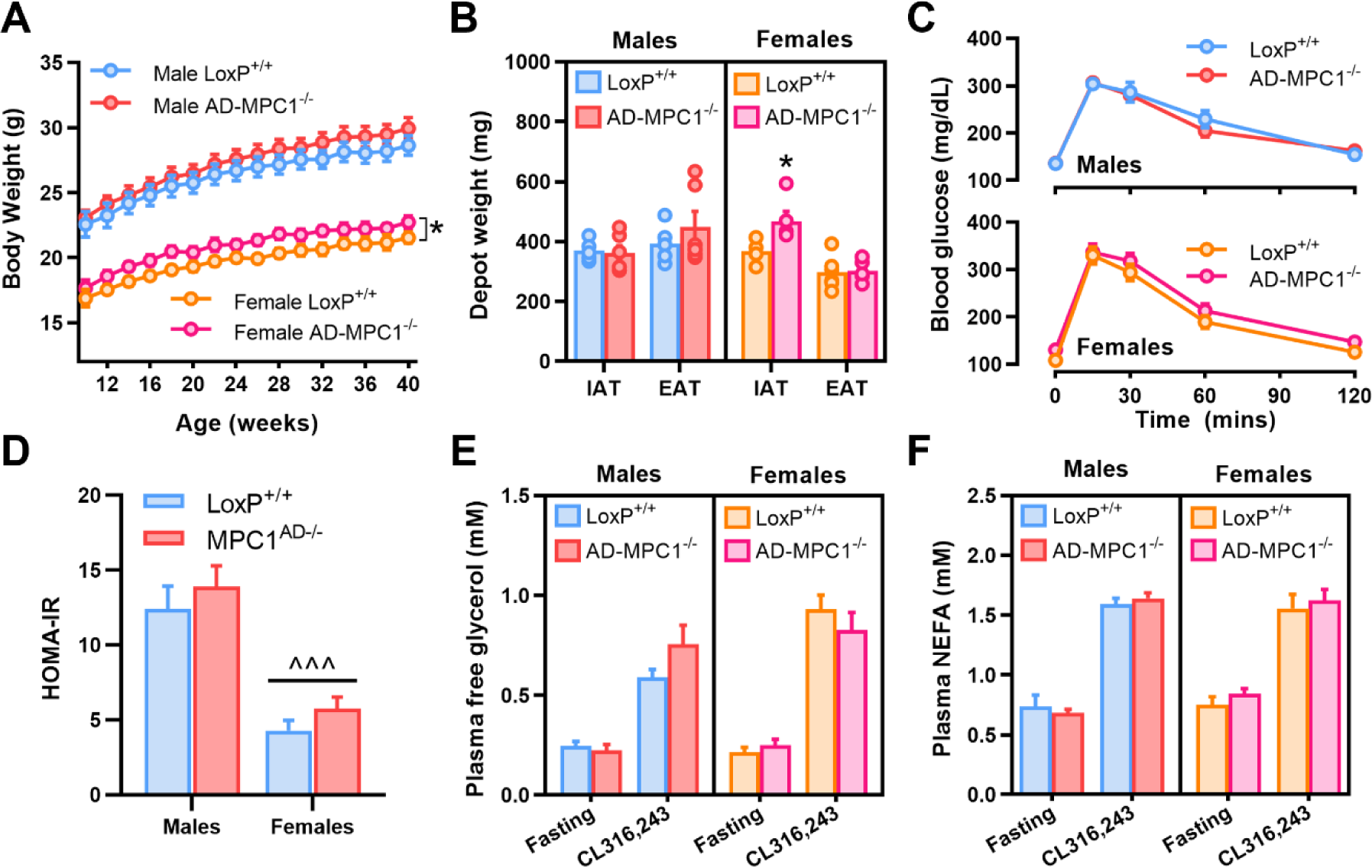
Sex-specific dependencies on the MPC for adipose glycerol-3-phosphate synthesis in vivo. (**A - D**) *In vivo* incorporation of intraperitoneally administered [U-^14^C] glucose into the glycerol moieties of total lipids (glyceroneogenesis **A**, **B**, **D**) or polar metabolites (**C**) in inguinal adipose (**A**), epididymal adipose (**B - C**), or liver (**D**) for male and female MPC^AD-/-^ mice or LoxP^+/+^ controls fed either a high fat western-style diet (WD) or a zero fat, sucrose enriched diet (ZFD) for 24 weeks. (**E** – **F**) mRNA expression of genes involved in glycerol metabolism in inguinal (**E**) or epididymal (**F**) adipose tissue for male and female MPC^AD-/-^ mice or LoxP^+/+^ controls fed a zero fat, sucrose enriched diet (ZFD). **P<0.01, ***P<0.001 for MPC^AD-/-^ vs LoxP^+/+^; ^#^P<0.05, ^###^P<0.001 main effect of genotype. Data are mean ± SE for 8-10 mice per group.

### Total adiposity is maintained in Mpc1^AD-/-^ mice fed a lipid restricted diet

As with chow fed mice, no overt metabolic phenotypes were evident when mice were fed either a WD or ZFD, with weight gain (Figure 8A and D and Supplemental Fig. 5A and C), body composition (Figure 8B and E) and glucose tolerance (Figure 8C and F and Supplemental Fig. 5B and D) comparable between *Mpc1*^AD-/-^ mice and LoxP^+/+^ controls. Thus, despite intrinsic impairments in pyruvate driven adipose DNL (Figure 3B and Supplemental Fig. 3A) and sex-specific impairments in glucose-mediated triglyceride assembly under conditions of dietary lipid restriction (Figure 4A-B and Figure 5A-B), total body fat accretion was entirely preserved in *Mpc1*^AD-/-^ mice regardless of sex or diet.

**Figure 8:**
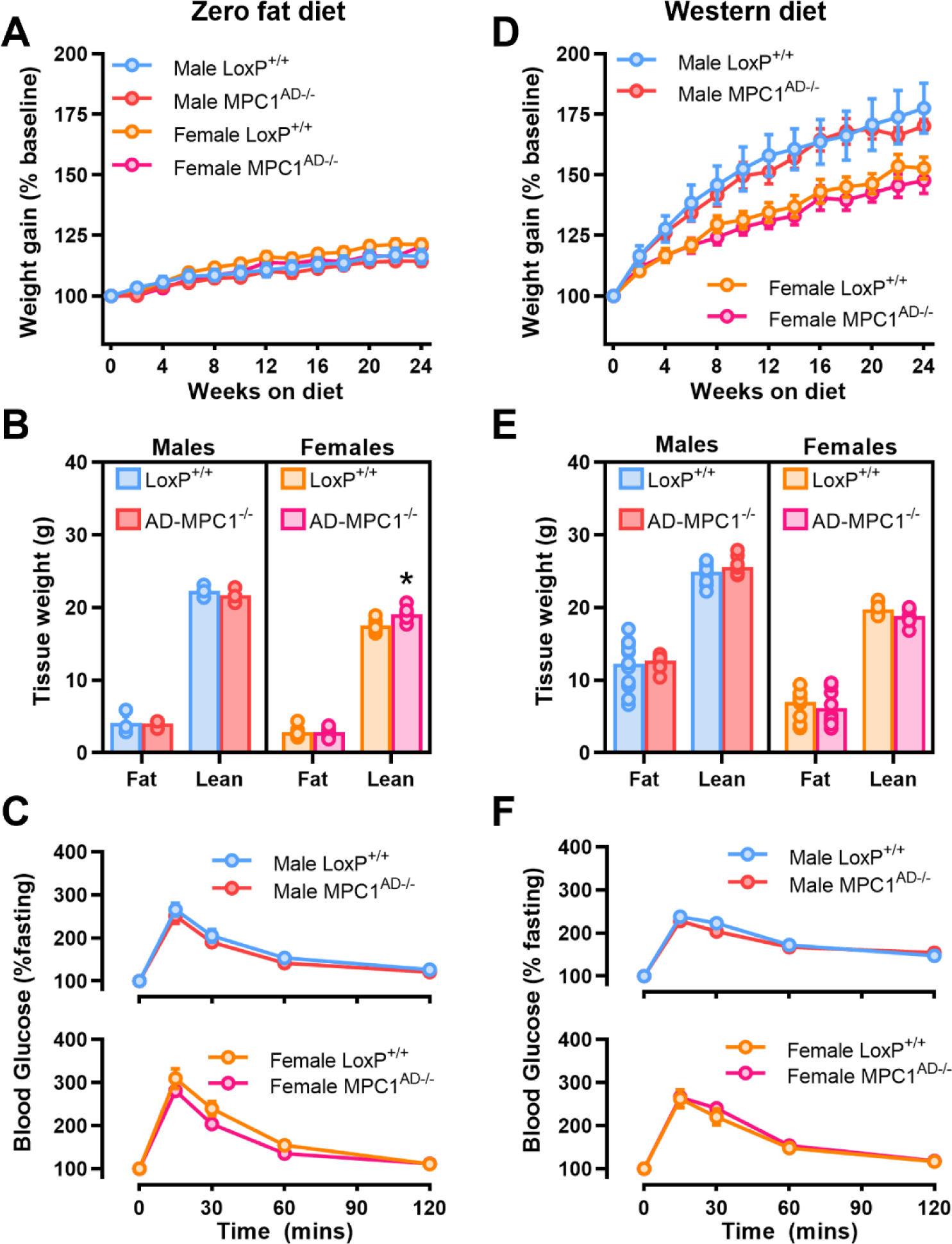
Decreased adipose TCA cycle pool size in female Mpc1AD-/- mice fed a lipid restricted diet. Polar metabolite abundances in epididymal adipose tissue from male and female MPC^AD-/-^ mice or LoxP^+/+^ controls fed a zero fat, sucrose enriched diet (ZFD) determined by gas chromatography mass spectrometry. (**A**) Summed and (**B**) individual TCA cycle intermediates, (**C**) alanine, (**D**) glutamate, (**E**) glutamine, (**F**) branched chain alpha keto acids, (**G**) the ratio of 3-hydroxybutyrate to alpha keto isovalerate and (**H**) glycerol-3-phosphate. Abundance data represent relative abundances of metabolites (normalized to labeled valine internal standard) except for alanine and glutamate (pmol / mg tissue). ^P<0.05, ^^P<0.01 for females vs males; *P<0.05, **P<0.01 main effect of genotype. Data are mean ± SE for 7-8 mice per group.

## DISCUSSION

Sexual dimorphism in adipose biology is believed to contribute to differential risk profiles for cardiometabolic diseases in males versus females, yet the molecular mechanisms underpinning these differences remain unclear. Here we reveal a sex-dependent role for the mitochondrial pyruvate carrier in adipose tissue glucose metabolism. Adipocyte-specific MPC depletion compromises the synthesis of fatty acids and glycerol-3-phosphate from glucose in female mice, whereas glucose trafficking into adipose triglycerides is entirely preserved in *Mpc1*^AD-/-^ males. Notably, these alterations in adipose glucose metabolism were only exposed when female mice were fed a carbohydrate rich, lipid restricted diet, consistent with an increased reliance on carbohydrates for endogenous triglyceride synthesis under these conditions (Aarsland et al., 1997). However, loss of adipocyte MPC did not impair body fat accumulation during dietary lipid restriction, regardless of sex. Instead, our findings support a compensatory role for MPC-independent pathways in sustaining triglyceride assembly and storage from non-glucose precursors in *Mpc1*^AD-/-^ female mice.

Inefficient deposition of dietary nutrients in adipose triglycerides is a purported pathophysiological feature of human obesity and insulin resistance (Allister et al., 2015). Whereas research in the obesity field has typically focused on the relevance of mitochondrial content and respiratory capacity, recent findings also highlight the involvement of adipocyte mitochondria in the regulation of nutrient storage (Zhu et al., 2022). Consistent with the central role of mitochondrial pyruvate transport in intermediary metabolism (Nathaniel 2014), our *in vitro* and *ex vivo* experiments confirmed that pharmacological blockade or genetic depletion of the MPC impairs the *de novo* synthesis of fatty acids and glycerol-3-phosphate from pyruvate in 3T3L1 adipocytes, whilst these pathways are almost completely abolished in mature adipose tissue explants from *Mpc1*^AD-/-^ mice. Altogether, these experiments indicate that at least one third of triglyceride accumulation in adipocytes depends upon MPC-gated metabolism, with the remainder presumably supported by alternate substrates which bypass the MPC (Gray et al., 2015; McCommis et al., 2015). Despite confirming an obligate role of the MPC for carbohydrate storage in adipose explants, *Mpc1*^AD-/-^ mice maintained comparable fat mass to control mice, even under conditions of dietary lipid restriction. This discrepancy suggests that in absence of MPC mediated DNL in adipose, whole body lipid synthesis is sustained either through i) MPC-independent DNL within adipocytes or ii) from non-adipose (e.g. hepatic) DNL. As will be discussed, our findings imply that the precise mechanism(s) may depend upon biological sex.

Carbohydrate flux into adipose triglycerides, whether measured *ex vivo* (from pyruvate) or *in vivo* (from glucose), was consistently greater in female adipose tissue than males. This was further supported by higher lipogenic gene expression in female adipose and parallels previous findings of higher rates of glucose storage in female adipose compared to males (Fernandez et al., 2019; Macotela et al., 2009). This higher capacity for adipose DNL could reflect an increased reliance on adipose tissue for whole body lipid synthesis in females, whereas males may obtain a greater contribution from non-adipose (i.e. liver) lipogenesis. This concept was previously illustrated in ZFD-fed mice deficient for adipose ATP citrate lyase (ACLY) (Fernandez et al., 2019) and is further supported by the current study. Thus, in the context of a relatively low requirement for adipose DNL, it is perhaps unsurprising that MPC-gated metabolism is dispensable for adipose triglyceride synthesis in male mice. In contrast to males, female *Mpc1*^AD-/-^ mice displayed intrinsic alterations in adipose metabolism, characterized by impaired nutrient storage pathways, reductions in TCA cycle intermediate pool sizes, and compensatory changes in lipogenic gene expression. Together, these data suggest that mitochondrial pyruvate transport may only be rate limiting for higher rates of DNL or gly-3P synthesis, including those observed in females consuming a ZFD.

A key phenotypic difference between female mice lacking adipose MPC and those lacking adipose ACLY is that, when consuming ZFD, *Mpc1*^AD-/-^ females did not become lipodystrophic nor develop any overt systemic metabolic abnormalities compared to littermate controls. Thus, whereas adipose ACLY appears conditionally essential for body fat accretion in females (Fernandez et al., 2019), reliance on MPC-gated metabolism for adipose accumulation may be obviated by compensatory pathways supporting adipose DNL from non-glucose precursors. Indeed, prior work has demonstrated considerable redundancy in the regulation of metabolic processes by the MPC (Gray et al., 2015; McCommis et al., 2015; Yiew et al., 2023). For example, cytosolic conversion of pyruvate to alanine circumvents mitochondrial pyruvate transport to support hepatic gluconeogenesis (McCommis et al., 2015). Interestingly, ZFD-fed *Mpc1*^AD-/-^ females displayed reduced adipose alanine concentrations, suggesting increased alanine utilization under these conditions. However, succinate and malate concentrations were also reduced which, consistent with MPC inhibition in hepatocytes (Yiew et al., 2023), underscores the obligate dependence on the direct import of mitochondrial pyruvate via the MPC to maintain TCA cycle intermediate pool sizes. Notably, lipogenic gene expression was increased in *Mpc1*^AD-/-^ female adipose, suggesting transcriptional compensation for the loss of glucose-derived lipogenic acetyl-CoA. Further experiments should consider whether cytosolic acetyl-CoA provision (e.g. from acetate) directly sustains adipose DNL in the face of MPC blockade in female adipose.

In addition to supporting adipose DNL, pyruvate is an important substrate for the *de novo* synthesis of gly-3P, required for acylglycerol assembly and accumulation. Glyceroneogenesis shares its initial enzymatic steps with the gluconeogenic pathway, diverging beyond the metabolism of dihydroxyacetone phosphate. Here we confirmed that genetic or pharmacological blockade of the MPC, like its role in pyruvate mediated gluconeogenesis in the liver (Gray et al., 2015; McCommis et al., 2015), impairs glyceroneogenesis from pyruvate in adipose tissue. Interestingly, the incorporation of circulating glucose into the glycerol fraction of acylglycerols was also impaired in MPC deficient adipose *in vivo*, but only in *Mpc1*^AD-/-^ females fed a ZFD. These findings mirror the partial suppression of glucose synthesis from glycerol via the indirect pathway (i.e. gluconeogenesis) in MPC deficient livers (Yiew et al., 2023). Our tracer methodology cannot discriminate between the direct and indirect synthesis of glycerol-3-phopshate from glucose. However, based upon the comparable indices of adipose glucose uptake between *Mpc1*^AD-/-^ and LoxP^+/+^ mice (Figure 7C), together with the finding that levels of free gly-3P in adipose were similar across genotypes (Figure 8H), it is likely that the direct pathway of gly-3P synthesis from glucose was preserved in *Mpc1*^AD-/-^ mice. This implies that adipose glyceroneogenesis was specifically compromised in *Mpc1*^AD-/-^ females fed a ZFD. Nevertheless, the maintenance of fat mass in *Mpc1*^AD-/-^ females illustrates that, even under lipid restricted conditions, gly-3P generation from glucose was not limiting to triglyceride accumulation. Our transcriptional data hint towards the upregulation of glycerol kinase as a potential compensatory mechanism, but other alternate pathways of gly-3P synthesis could also contribute.

It should be acknowledged that tracing pathways of [U-^14^C] glucose *in vivo* is complicated by the potential recycling of tracer between tissues (e.g. liver, skeletal muscle). However, based upon prior experiments, the contribution of non-adipose tracer metabolism to circulating labeled metabolites is expected to have a relatively minor influence on the measurement of adipose DNL or glycerol-3-phosphate synthesis over the time course of the studies performed here (Björntorp et al., 1970).

In conclusion, we demonstrate that the mitochondrial pyruvate carrier facilitates the increased trafficking of glucose into both fatty acids and gly-3P in female adipose tissue under lipid restricted conditions. Disruption of this process does not alter the whole-body phenotypic adaptation to dietary stress, highlighting the remarkable flexibility for mitochondrial substrate selection towards metabolic homeostasis. Our findings are highly consistent with a recent report that loss of liver MPC constrains hepatic DNL during fasted refeeding in mice without impacting hepatic steatosis or circulating lipid concentrations (Yiew et al., 2023). However, our findings provide the first evidence of sexual dimorphism in MPC-gated metabolism and thus advance understanding of the molecular mechanisms underlying sex differences in adipose biology and metabolic physiology.

## ACKNOWLEDGEMENTS

C.E.S. is supported by funding from the European Union’s Horizon 2020 research and innovation programme under Marie Skłodowska-Curie grant agreement No 945425 and by a Research and Knowledge Exchange Award from The Physiology Society (Grant number TPSRKE02). L.N. is supported by funding from the NIDDK (R01DK128247). The polar metabolomics data was generated with valuable support from Dr Xiaofei Yin and Professor Lorraine Brennan within the UCD Conway Institute of Biomolecular and Biomedical Research Metabolomic Facility.

## AUTHOR CONTRIBUTIONS

Conceptualization and experimental design: C.E.S. and L.N. Methodology: C.E.S., T.B., and M.J.F. performed the cell culture experiments, animal experiments and biochemical analyses; C.E.S. and M.W. performed the metabolite analyses. All authors contributed to the data generation. Writing: C.E.S. prepared figures and drafted the original manuscript; All authors reviewed and approved the final manuscript draft. Funding acquisition: C.E.S. and L.N.

## SUPPLEMENTAL FIGURE LEGENDS

**<u>Supplemental Figure 1 (related to Figure 1):</u> siRNA mediated depletion of MPC1 in 3T3L1 adipocytes**

Representative western blot and normalized, quantified protein expression of the mitochondrial carrier protein 1 (MPC1) in differentiated 3T3L1 adipocytes following reverse transfection with MPC1 siRNA. Cells were reverse transfected at 5 days post-differentiation as described in materials and methods. MPC1 protein expression was quantified in cell lysates at 2-, 3-, 5-, and 8-days post-transfection to determine the optimal time for subsequent assays in transfected cells (shown in Figure 1E).

**<u>Supplemental Figure 2 (related to Figure 2):</u> Metabolic phenotype of human cohorts**

(**A**) Blood glucose, (**B**) plasma insulin, and (**C**) plasma free fatty acid concentrations during an oral glucose tolerance test, as well as the incremental area under (AUC) the glucose x time curve (**D**), the ratio of the glucose AUC / insulin AUC (**E**), and the adipose insulin resistance index (**F**) in human subjects with normal glucose tolerance (NGT) or with combined impaired fasting glucose and impaired glucose tolerance (IFG/IGT). *P<0.05, **P<0.01 for NGT vs IFG/IGT. Data are n=7 in each group.

**<u>Supplemental Figure 3 (related to Figure 3):</u> Further characterization of adipose explants from MPC^AD-/-^ mice**

(**A**) *Ex vivo* incorporation of [2-^14^C] pyruvate into the fatty acyl moieties of total lipids in inguinal adipose explants from male and female MPC^AD-/-^ mice or LoxP^+/+^ controls. (**B - E**) Rates of glycerol release (**B**), and non-esterified fatty acid release (**D**, **E**), and calculated non-esterified fatty acid re-esterification (**C**) from epididymal (EAT) or inguinal (IAT) adipose explants from male MPC^AD-/-^ mice or LoxP^+/+^ controls under basal (unstimulated) conditions or during treatment with insulin or forskolin plus triacsin C. *P<0.05, **P<0.01 for MPC^AD-/-^ vs LoxP^+/+^. Data are mean ± SE for at least five mice per group.

**<u>Supplemental Figure 4 (related to Figure 5):</u> Further characterization of MPC^AD-/-^ mice under dietary stress**

(**A** - **B**) Absolute body weight and (**C** – **D**) blood glucose concentrations during an oral glucose tolerance test for male and female MPC^AD-/-^ mice or LoxP^+/+^ controls fed either a zero fat, sucrose enriched diet (ZFD **A - B**) or a high fat western-style diet (WD **C - D**) for 24 weeks. **P<0.01 for MPC^AD-/-^ vs LoxP^+/+^. Data are mean ± SE for 8-10 mice per group.

**<u>Supplemental Figure 5 (related to Figure 6):</u> Further characterization of lipogenic gene expression and liver contribution**

(**A - B**) mRNA expression of lipogenic genes in inguinal adipose tissue iWAT, (**C**) *in vivo* incorporation of intraperitoneally administered [U-^14^C] glucose into the fatty acyl moieties of total lipids (lipogenesis) and (**D**) mRNA expression of lipogenic genes in livers from male and female MPC^AD-/-^ mice or LoxP^+/+^ controls fed either a high fat western-style diet (WD) or a zero fat, sucrose enriched diet (ZFD) for 24 weeks. ^P<0.05 for female vs male. Data are mean ± SE for 8-10 mice per group.

