## Supplemental figures for "Sex-dependent adipose glucose partitioning by the mitochondrial pyruvate carrier"

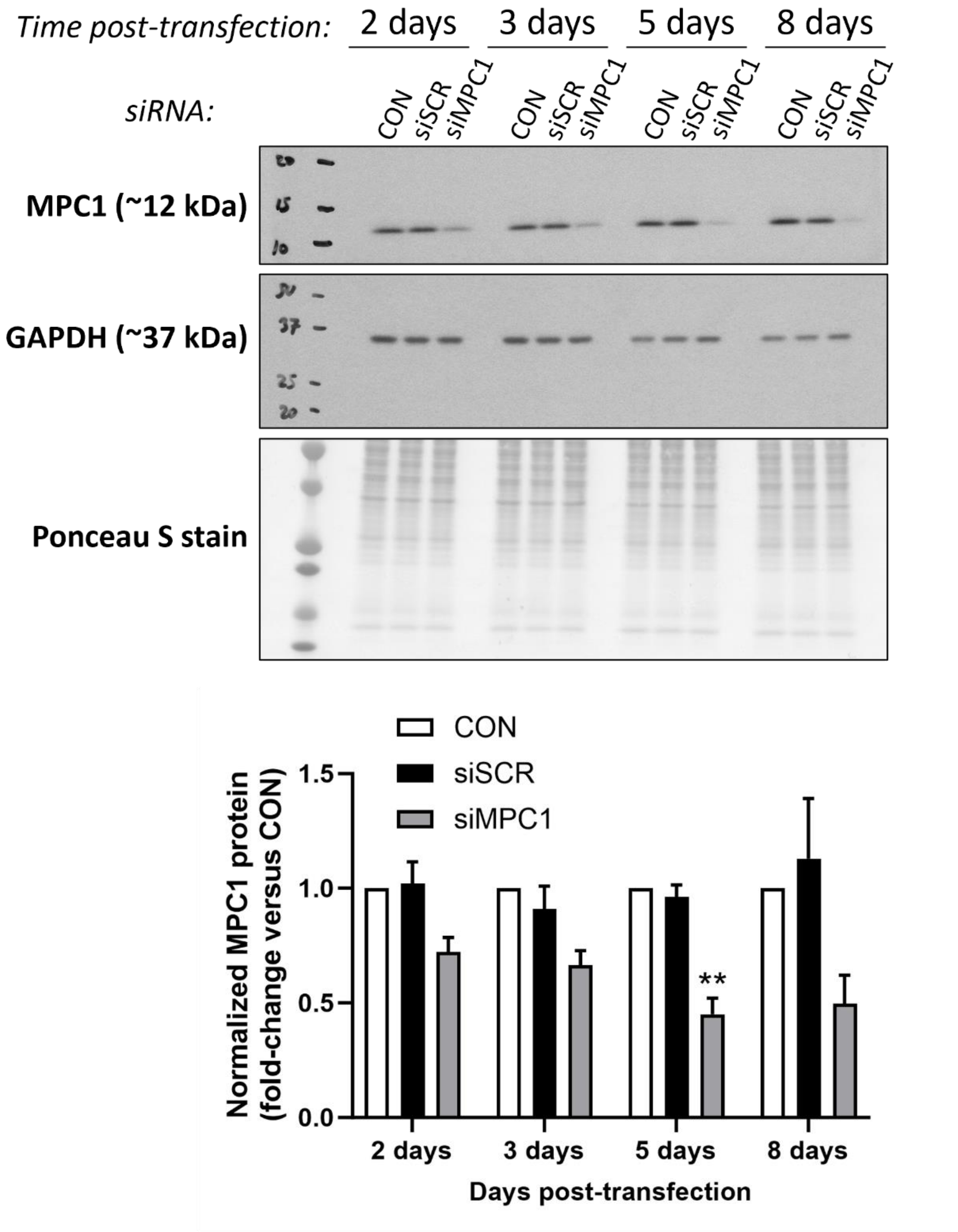


**Supplemental Figure 1**

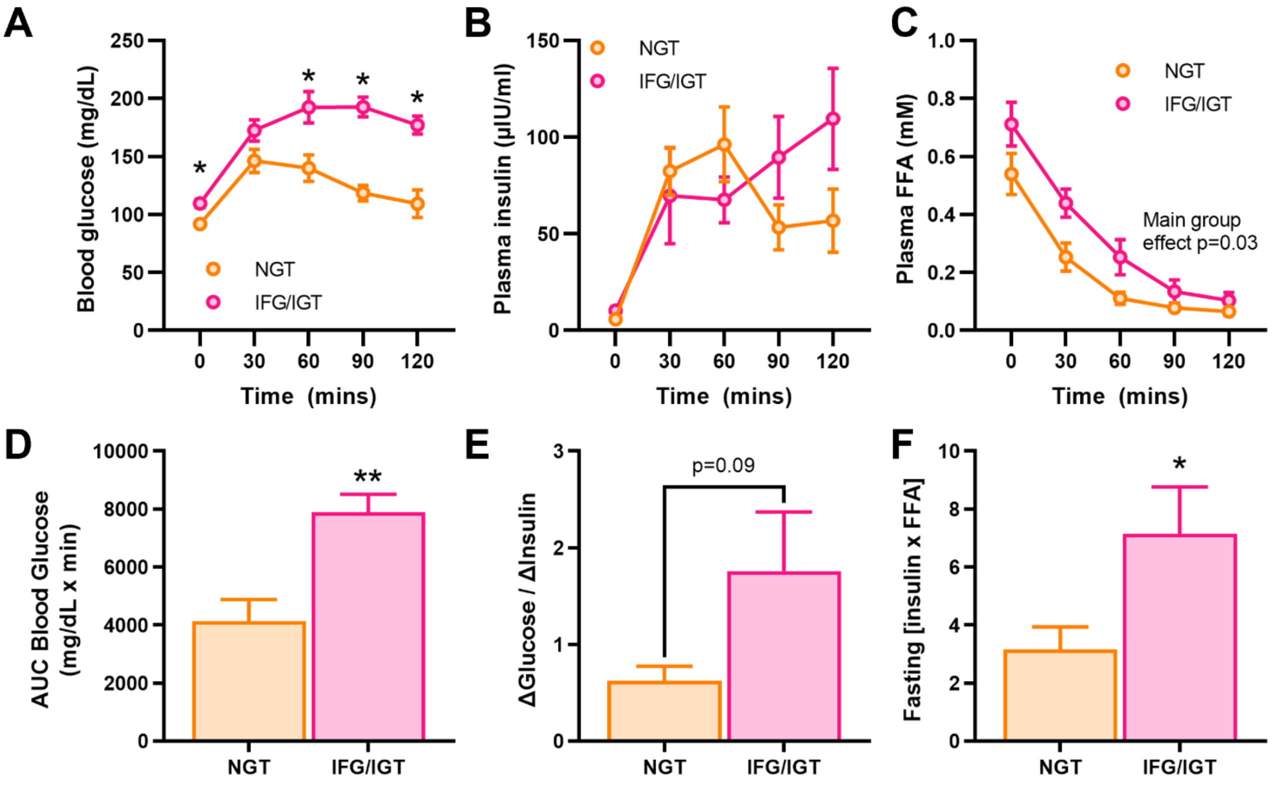


**Supplemental Figure 2**

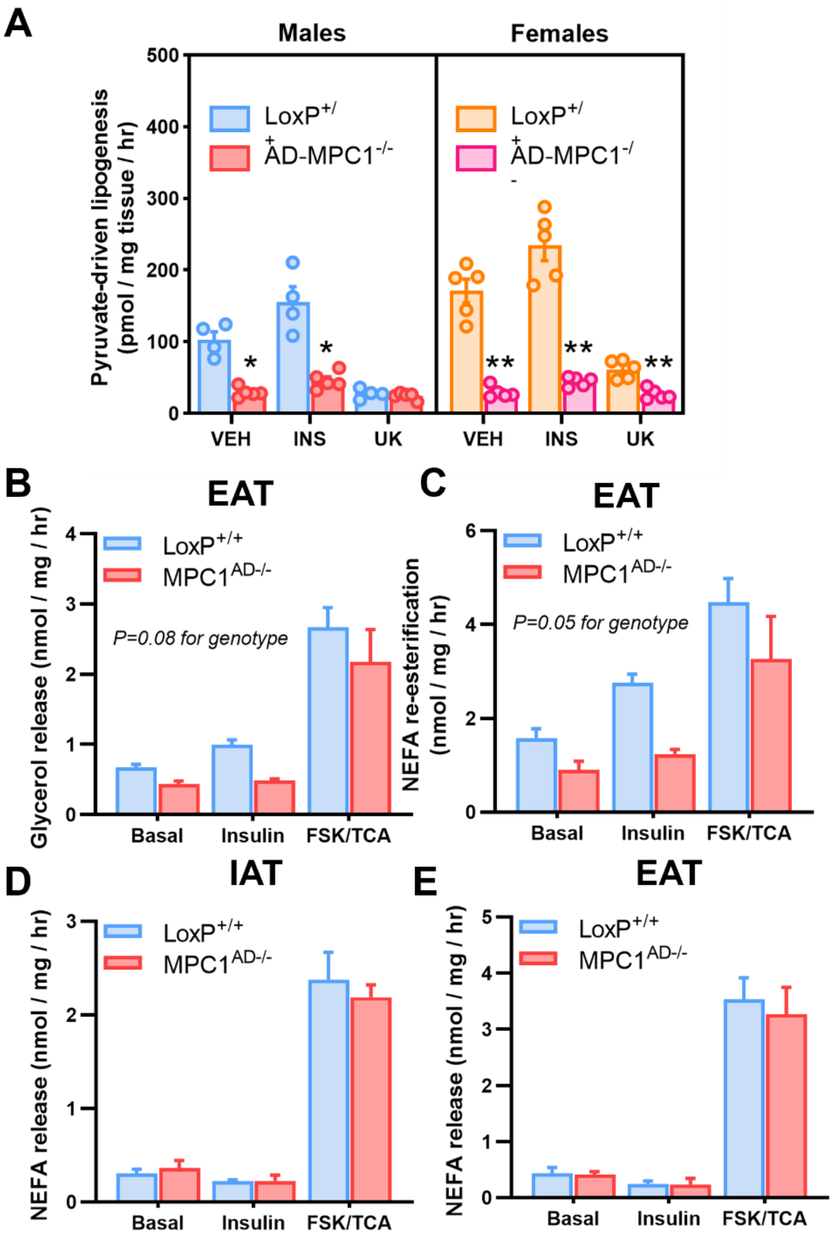


**Supplemental Figure 3**

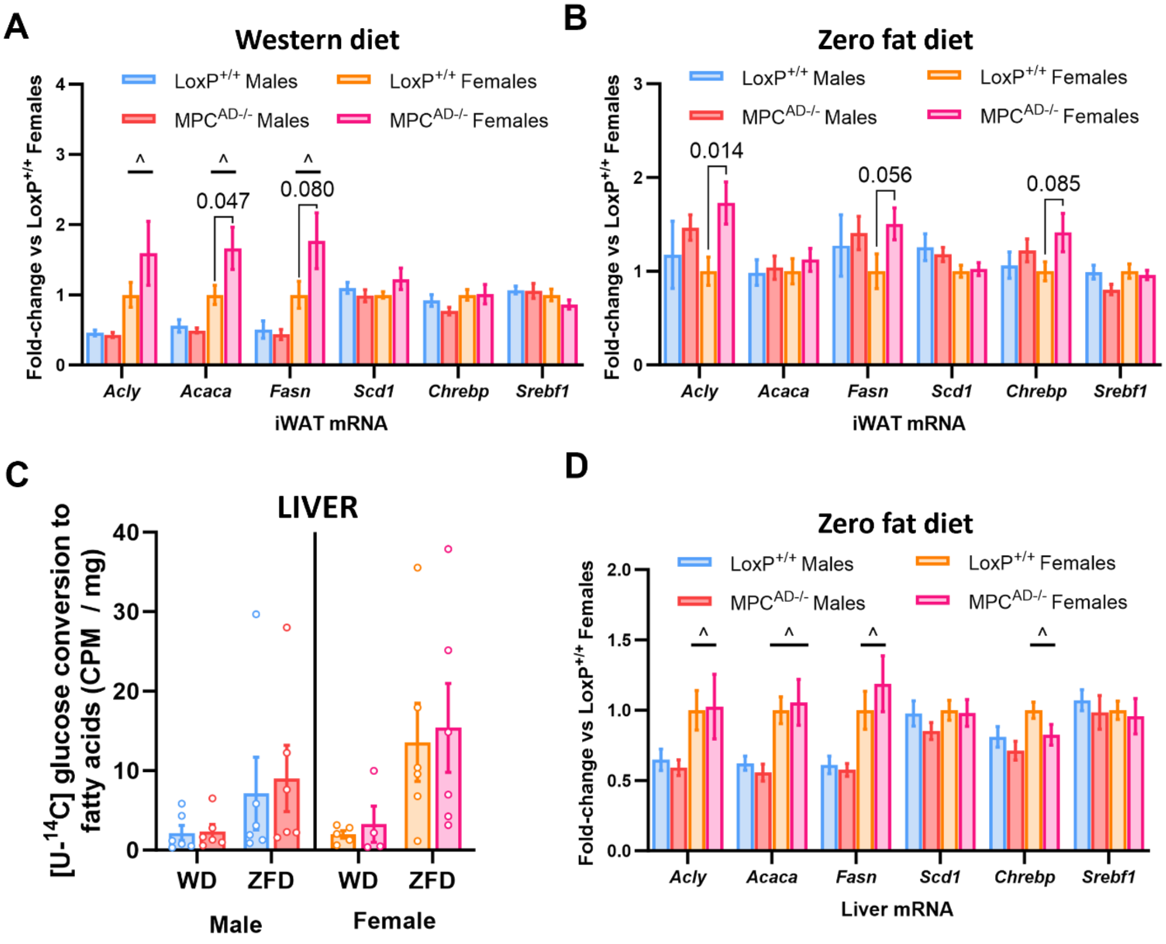


**Supplemental Figure 4**

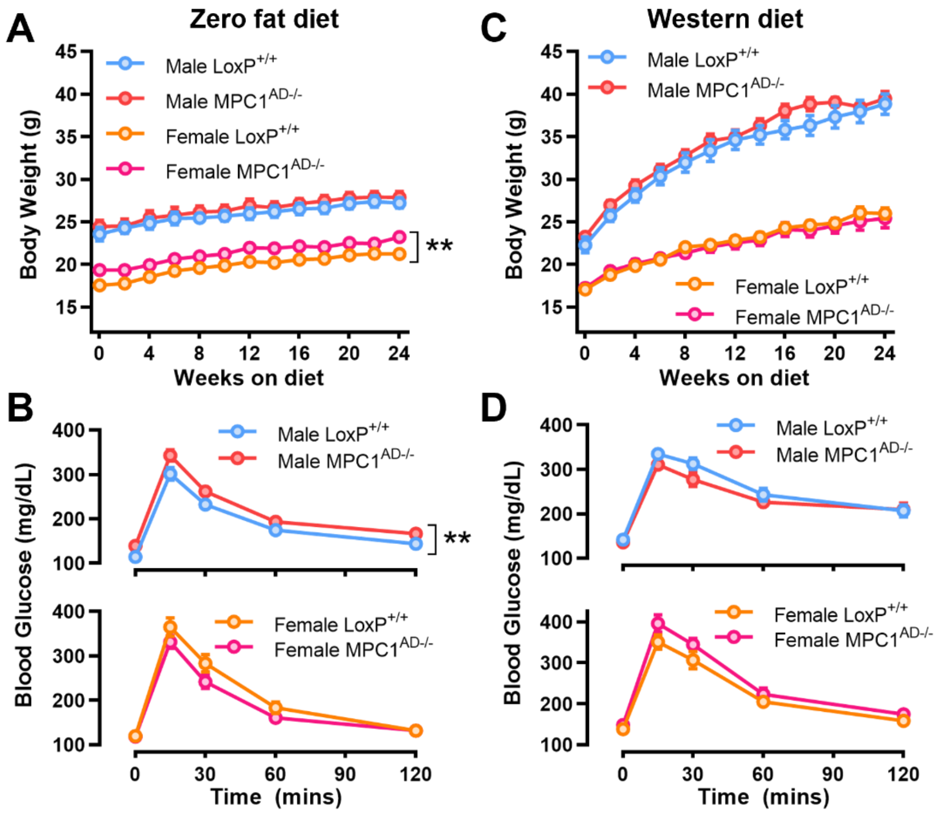


**Supplemental Figure 5**
